# Hypothalamic control of noradrenergic neurons stabilizes sleep

**DOI:** 10.1101/2023.10.22.563502

**Authors:** Gianandrea Broglia, Giorgio Corsi, Pierre-Hugues Prouvot Bouvier, Anne Vassalli, Mehdi Tafti, Mojtaba Bandarabadi

**Affiliations:** Department of Biomedical Sciences, University of Lausanne, Lausanne, Switzerland

## Abstract

Hypocretin/orexin neurons are essential to stabilize sleep, but the underlying mechanisms remain elusive. We report that hypocretin neurons of the perifornical hypothalamus are highly active during rapid eye movement sleep and show state-specific correlation with noradrenergic neurons. Deletion of hypocretin gene significantly increased periodic reactivations of locus coeruleus noradrenergic neurons during sleep and dysregulated their activity across transitions, suggesting a role for hypocretin neurons in mediating neuromodulation to stabilize sleep.

## Main text

Spontaneous transitions in vigilance states have deterministic neural substrates allowing the maintenance of stable sleep-wake cycle over time^1^. Complex interconnected wake-promoting and sleep-generating pathways mutually inhibit each other to induce rapid and complete state transitions^2,3^. Hypocretin/orexin (HCRT) neurons are critical for the proper regulation of these transitions^4^, in addition to attention, motivation, and appetite^5,6^. Hypocretin neurons are localized in the lateral hypothalamus (LH), perifornical area of hypothalamus (PeF), and a sparse distribution in the dorsomedial hypothalamus (DMH)^7,8^. They project to all wake-promoting monoaminergic and cholinergic nuclei, the midline thalamus, and the cortex^9^, and can promote wakefulness and maintain arousal by stimulating these areas. Deficiency in hypocretin neurotransmission causes the sleep disorder narcolepsy, associated with chronic sleepiness, fragmented sleep, abnormal transitions to rapid eye movement sleep (REMS), and sudden loss of muscle tone during wakefulness in both humans and animals^10–12^.

Locus coeruleus noradrenergic (LC^NA^) neurons, which receive the densest projections of hypocretin cells^9^, show periodic reactivations during non-rapid eye movement sleep (NREMS) and can gate sleep to wake transitions^13,14^. Optogenetic activation of LH^HCRT^ or LC^NA^ neurons induces rapid awaking from both NREMS and REMS^15,16^. Hypocretin neurons play a crucial role in stabilizing vigilance states in the current theoretical models, e.g. the flip-flop switch model^3,17^. However, LH^HCRT^ neurons are relatively silent during sleep^18^, and the current models cannot explain long-term regulation of sleep-wake cycle in a steady manner in the absence of hypocretin^19,20^. Therefore, other hypocretin subpopulations or mechanisms might be involved to maintain consolidated sleep and regulate vigilance state transitions.

First, to test whether PeF^HCRT^ neurons have different temporal dynamics from LH^HCRT^ cells across sleep-wake states and during spontaneous transitions, we performed *in vivo* calcium imaging in the PeF and LH, combined with EEG/EMG recordings in freely behaving *Hcrt-IRES-Cre* mice (Supp. Fig. 1,2). While LH^HCRT^ cells are mainly active during wakefulness, as in previous reports^18,21,22^, we unexpectedly found that PeF^HCRT^ neurons are highly active during REMS and wakefulness (Fig. 1a-c). We also recorded a weak calcium signal from LH^HCRT^ neurons during REMS (Fig. 1b,c), consistent with a recent study reporting that a small subpopulation of LH^HCRT^ neurons are active during REMS^22^. Both PeF^HCRT^ and LH^HCRT^ neurons also showed sparse activity during NREMS, with a peak at NREMS to wake transitions (Fig. 1b). To investigate the correlation between activity of PeF^HCRT^ neurons and the noradrenergic system, we simultaneously recorded PeF^HCRT^ calcium signal and regional norepinephrine release in the LC and in its major projecting target of the paraventricular nucleus of the thalamus (PVT). We found that PeF^HCRT^ neuronal activity correlates with norepinephrine release in both LC and PVT during wakefulness and NREMS, while anti-correlated during REMS (Fig. 1d), suggesting different PeF^HCRT^ subpopulations that are active during wake/NREMS and REMS.

**Fig. 1:**
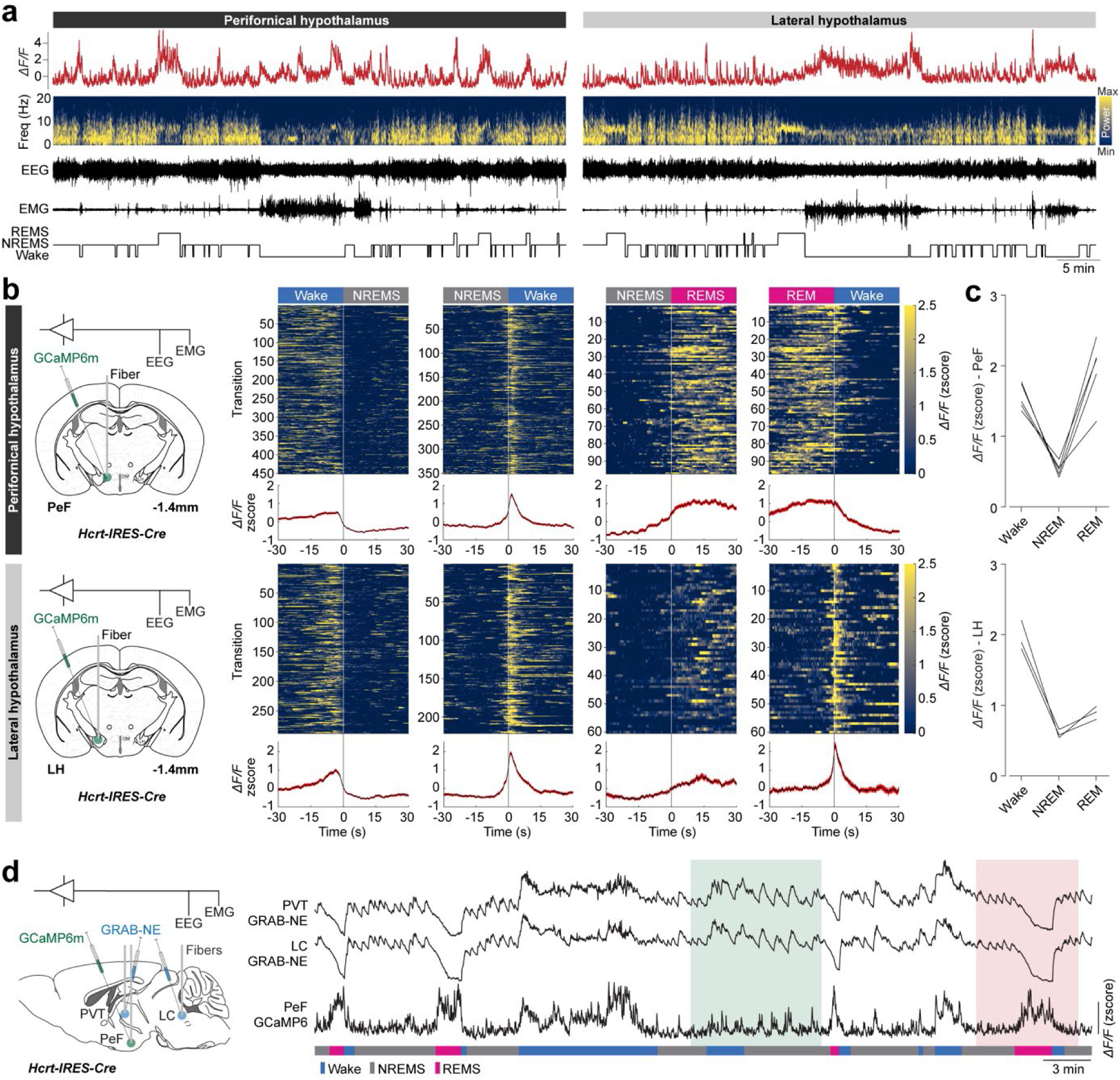
Hypocretin neurons of the perifornical hypothalamus are REMS-active. **a)** Representative fiber photometry calcium activity of hypocretin neurons in the PeF and LH, time-frequency representation of EEG signals, and raw EEG/EMG signals. Hypnogram is depicted below. **b)** Schematic coronal sections with fiber location for each condition, and transition panels of vigilance states for the PeF and the LH hypocretin neurons. Heatmaps show the normalized color-coded *ΔF/F* for each transition (top) and traces represent the averaged signals for the 30 sec before and after the transitions (bottom). Data represent recordings from 5 mice for the PeF and 3 mice for the LH. **c)** Average *ΔF/F* of calcium signal across each vigilance state during the whole recording session (12:00-15:00, ZT4-7; n = 5/3 mice for the PeF/LH). **d)** State-specific correlation between activity of PeF hypocretin neurons and noradrenergic activity. Representative simultaneous multisite fiber photometry recordings of PeF hypocretin neurons using GCaMP6m and regional norepinephrine release in the LC and PVT using GRAB_NE_. Green box highlights NREMS and wake activity and red one a REMS episode.

Next, to investigate the activity of LC^NA^ neurons in the absence of hypocretin, we generated hypocretin knockout (*Hcrt^KO/KO^*) mice harboring Cre recombinase in noradrenergic neurons by breeding *Hcrt^KO/KO^* and *Dbh-Cre* mice (*Hcrt^KO/KO^*x*Dbh-Cre*). We then performed *in vivo* calcium imaging of LC^NA^ neurons in freely behaving knockout (*Hcrt^KO/KO^*x*Dbh-Cre*) and control (*Dbh-Cre*) mice, combined with EEG/EMG recordings. We found that LC^NA^ neurons are periodically reactivated during NREMS, as reported recently^13,14^, in both genotypes (Fig. 2a). Multisite fiber photometry of the LC and its projections in the PVT and cingulate cortex revealed that the norepinephrine release during these reactivations reaches both thalamic and cortical sites, similar as during wakefulness (Supp. Fig. 3). LC^NA^ neuronal activity increased prior to NREMS to wake transitions in both knockout and control mice (Fig. 2b), consistent with the role of LC^NA^ reactivations in providing a fragility window for transitions^13,14^. However, we found that LC^NA^ reactivations during NREMS are significantly more frequent in hypocretin knockout mice compared to controls (*p* = 0.0005, unpaired *t*-test), which resulted from a shorter refractory period (*p* < 0.0001, unpaired *t*-test; Fig. 2c). *Hcrt^KO/KO^* mice showed fragmented NREMS episodes, reflected in higher number of shorter bouts (number: *p* = 0.0017, duration: *p* = 0.0015, unpaired *t*-test; Fig. 2d), but similar total NREMS amount compared to controls (Supp. Fig. 4a,b). These data suggest a mediatory role for the hypocretin system during NREMS, where shorter sleep episodes result from hyperactivated LC^NA^ neurons in the absence of hypocretin signaling.

**Fig. 2:**
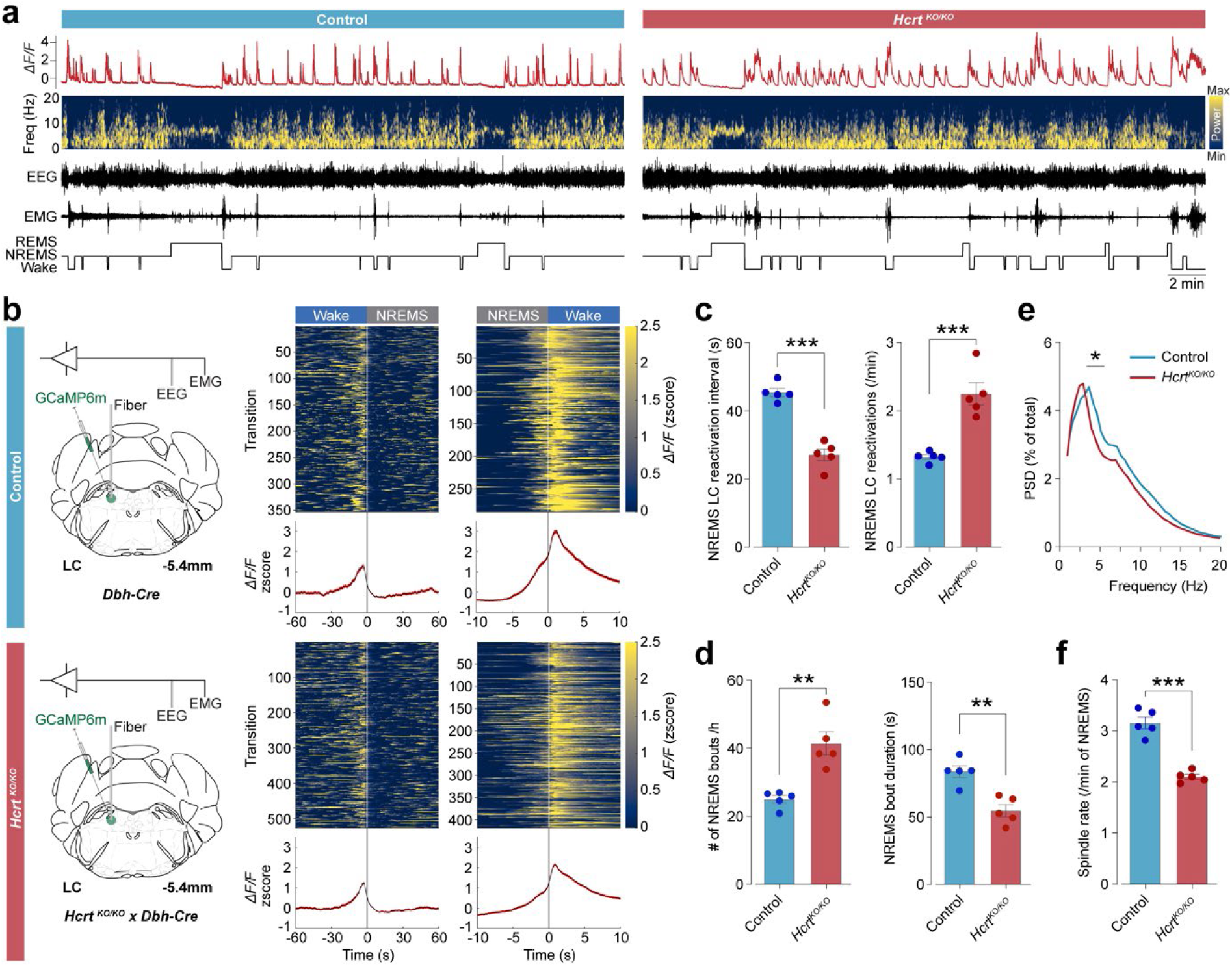
Hypocretin mediates LC noradrenergic reactivations during NREMS. **a)** Representative fiber photometry calcium activity of LC noradrenergic neurons, time-frequency representation of EEG signals, and raw EEG/EMG signals, from control and hypocretin knockout mice. Hypnogram is depicted below. **b)** Heatmaps show the normalized color-coded *ΔF/F* for each transition (top) and traces represent the averaged signals across the transitions (bottom). Data represent recordings from 5 control (*Dbh-Cre*) and 5 hypocretin knockout (*Hcrt^KO/KO^xDbh-Cre*) mice. **c)** Quantification of LC noradrenergic reactivations per minute of NREMS (*p* = 0.0005, unpaired *t*-test), and the time interval between LC reactivations during NREMS (*p* < 0.0001, unpaired *t*-test). **d)** Number of NREMS bouts per hour (12:00-15:00; ZT4-7) (*p* = 0.0017, unpaired *t*-test) and mean duration of NREMS bouts in the same recording session (*p* = 0.0015, unpaired *t*-test). **e)** PSD analysis during NREMS (two-way ANOVA, interaction: *p* < 0.0001, followed by Sidak test). **f)** number of spindles per minutes of NREMS (*p* < 0.0001, unpaired *t*-test). n = 5 mice per group.

We next investigated the effects of overactivity of LC^NA^ neurons in *Hcrt^KO/KO^* mice on the brain oscillations during NREMS. We calculated the power spectral density (PSD) of EEG signal and found a shift in the peak frequency of delta oscillations in *Hcrt^KO/KO^* mice compared to controls (Fig. 2e). As LC^NA^ activity is detrimental for the generation of thalamocortical sleep spindles^13^, we also quantified spindles and found a significant reduction in the NREMS spindle density and alternations in spindle characterizations (*p* < 0.0001, unpaired *t*-test; Fig. 2f and Supp. Fig. 5). These results indicate the contribution of subcortical neuromodulatory pathways in modulation of thalamocortical brain oscillations during NREMS.

Next, we tested how the absence of hypocretin signaling affects REMS architecture and its components. Although the LC^NA^ neurons are periodically reactivated during NREMS, the noradrenergic tone is completely absent tens of seconds prior to REMS onset and during REMS (Fig. 2a and Fig. 3a). However, we found that this silent period of LC^NA^ neurons prior to transitions to REMS is significantly shorter in *Hcrt^KO/KO^* mice compared to controls (*p* = 0.0279, unpaired *t*-test; Fig. 3a,b). Furthermore, LC^NA^ neuronal activity is very high immediately before REMS-to-wake transitions, but the lag between the activation of LC^NA^ neurons and transitions is significantly longer in *Hcrt^KO/KO^* mice compared to controls (*p* = 0.0077, unpaired *t*-test; Fig. 3a,c). *Hcrt^KO/KO^* mice also showed fragmented REMS episodes, reflected in higher number of shorter bouts (number: *p* < 0.0001, duration: *p* = 0.0162, unpaired *t*-test; Fig. 3d), and higher total REMS amount compared to controls (Supp. Fig. 4c,d). These results suggest that altered REMS architecture of *Hcrt^KO/KO^* mice may result from the dysregulated LC^NA^ activity during sleep.

**Fig. 3:**
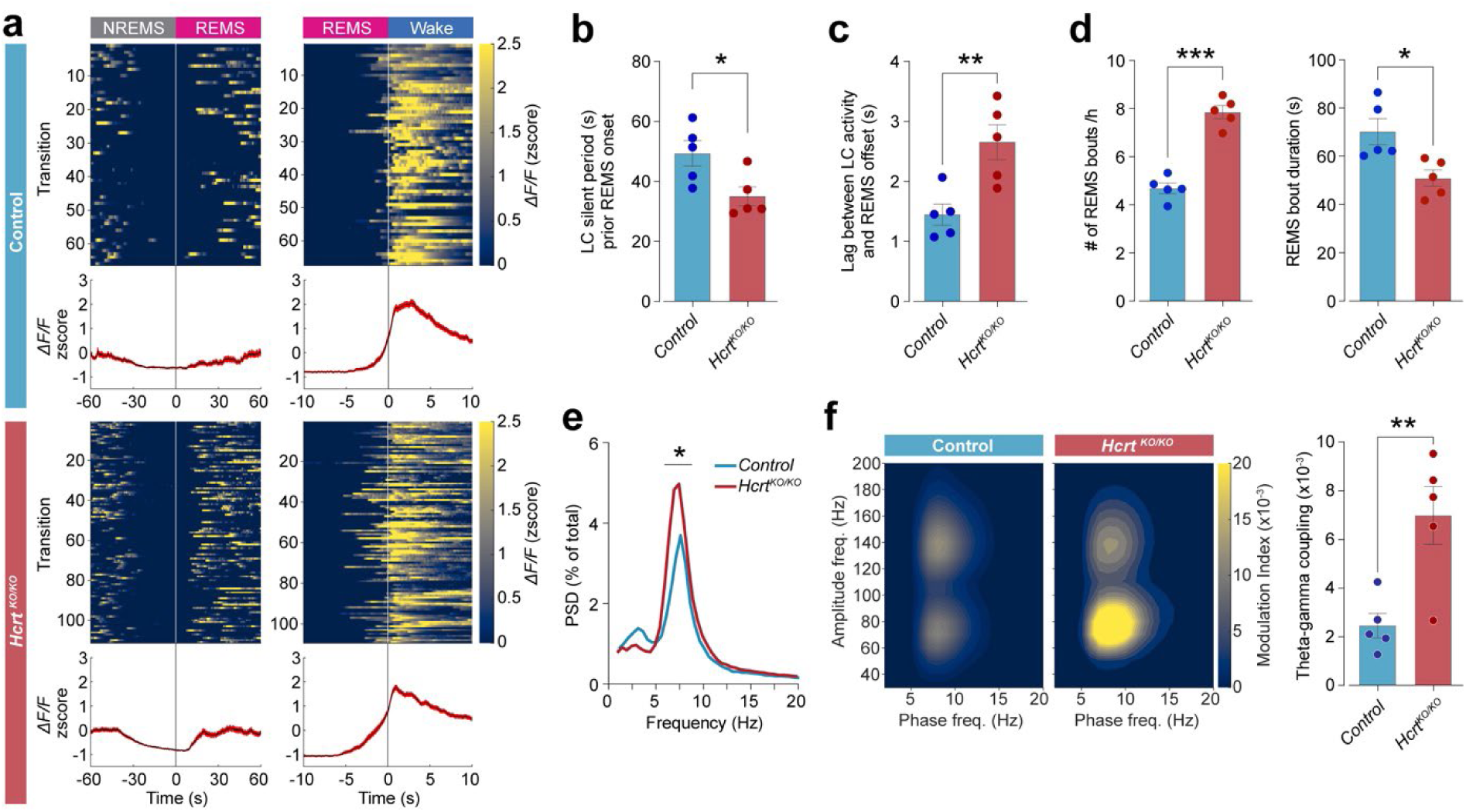
Lack of hypocretin dysregulates noradrenergic activity at REMS onset and offset. **a)** Transition panel for REMS onset and offset in *Dbh-Cre* (top) and *Hcrt^KO/KO^xDbh-Cre* mice (bottom). Normalized color-coded *ΔF/F* of photometry Ca^2+^ activity for individual transitions and averaged signal across all the transitions. Data represent transitions for 5 (*Dbh-Cre*) and 5 (*Hcrt^KO/KO^xDbh-Cre*) mice. **b)** LC silent period prior to REMS onset (*p* = 0.0279, unpaired *t*-test). **c)** Lag between LC noradrenergic activity and REMS termination (*p* = 0.0077, unpaired *t*-test). **d)** Number of REMS bouts per hour (12:00-15:00; ZT4-7) (*p* < 0.0001, unpaired *t*-test) and mean duration of REMS bouts in the same recording session (*p* = 0.0162, unpaired *t*-test). **e)** Power spectral density (PSD) during REMS (two-way ANOVA, interaction: *p* < 0.0001, followed by Sidak test). **f)** Heatmaps show comodulogram analysis for phase-amplitude cross-frequency coupling, and bars indicate the theta-gamma coupling level during REMS (*p* = 0.0078, unpaired *t*-test). n = 5 mice per group.

To assess the contribution of hypocretin neurons in the regulation of REMS oscillations, we calculated the PSD of EEG recordings and found an increased theta power in *Hcrt^KO/KO^* mice compared to controls (Fig.3e). We also investigated the interaction of theta rhythms with gamma oscillations, where theta phase modulates gamma power, and found significant increase in theta-gamma, but not theta-fast gamma, coupling in *Hcrt^KO/KO^* mice compared to controls (*p* = 0.0078, unpaired *t*-test; Fig.3f and Supp. Fig. 6a). These results suggest an inhibitory role for REMS-active hypocretin neurons in theta generation and modulation pathways. As previous reports showed that hypocretin neurons discharge during phasic REMS events^23,24^, we also quantified these events, but did not find alternations in the amount and characteristics of phasic REMS events between knockout and control animals (Supp. Fig. 6b), indicating only a correlation and not causation, between hypocretin activity and phasic REMS.

In summary, we found a dichotomy between the LH and PeF subpopulations of hypocretin neurons, where LH^HCRT^ neurons are principally wake-active, while PeF^HCRT^ neurons are mainly REMS-active. Although the role of LH^HCRT^ neurons in the regulation of arousal levels, reward, feeding, and emotional behaviors is well established^5,6,21^, the PeF/DMH neurons are hypothesized to regulate vigilance states only^25^. To our knowledge, our study is the first to report *in vivo* activity of PeF^HCRT^ neurons across vigilance states and during sleep-wake transitions and show that these neurons are highly active during sleep. Although we did not investigate the functional role of PeF^HCRT^ neurons directly, considering that these neurons are mainly REMS-active, they could be responsible for altered architecture and components of REMS in *Hcrt^KO/KO^* mice, e.g. the shorter epochs and higher theta power.

Deletion of hypocretin gene resulted in more frequent LC^NA^ reactivations during NREMS. As these reactivations are a window of sleep fragility and possible awakenings^13,14^, the fragmented NREMS in *Hcrt^KO/KO^* mice could be explained by a mediatory role for the hypocretin system that suppresses the LC^NA^ reactivations during sleep. Moreover, a shorter silent period of LC^NA^ neurons required prior to REMS onset can explain more frequent NREMS-REMS transitions in *Hcrt^KO/KO^* mice, and a similar mechanism could cause “sleep onset REM period” in human narcolepsy. Additionally, LC^NA^ activation prior to REMS termination appears earlier in *Hcrt^KO/KO^* mice compared to controls, suggesting a possible mechanism behind shorter REMS episodes and their premature termination, and indicating again a mediatory role for REMS-active hypocretin neurons in stabilization of REMS. Although the underlying mechanisms that drive periodic activation of the LC^NA^ neurons, and possibly other monoaminergic neurons, during NREMS are unclear, our results indicate that the hypocretin system has a major role in controlling the timing of this infra-slow oscillation. Altogether, our findings provide a mechanistic answer to the fragmented, but still regulated, sleep in the absence of hypocretin, as in human narcolepsy.

## Methods

### Animals

*Hcrt-IRES-Cre* knock-in (*Hcrt^tm1.1(cre)Ldl^*) heterozygote^26^ and *Dbh-Cre* (*Tg(Dbh-icre)1Gsc*) heterozygote^27,28^ mice were locally backcrossed into C57BL/6J mice. Homozygous hypocretin-knockout (*Hcrt^KO/KO^*) mice were used as a mouse model of narcolepsy^12^. *Hcrt^KO/KO^xDbh-Cre* mice were generated locally by crossing *Dbh-Cre* mice with *Hcrt^KO/KO^* mice to investigate noradrenergic neuronal activity in the absence of hypocretin. Genotyping of the lines was carried out using PCR. Animals at 12-18 weeks old at the time of experiments were group-housed with *ad libitum* access to standard food pellets and water at the constant temperature (23±1 °C) and humidity (30-40%), and a 12h/12h light/dark cycle with lights on from 8:00 a.m. to 8:00 p.m. All procedures were carried out in accordance with the Swiss federal laws and approved by the veterinary office of the State of Vaud, Switzerland.

### Stereotaxic viral injections

Adeno-associated viruses (AAV) carrying the fluorescent reporter were produced by the viral vector facility of University of Zurich, Switzerland (www.vvf.uzh.ch). To record neuronal activity, a Ca2+ sensor (ssAAV-9/2-hSyn1-chI-dlox-GCaMP6m(rev)-dlox-WPRE-SV40p(A), titer 4.2×10E12 vg/ml) was delivered in Cre-expressing areas of interest in the transgenic mice. For screening neurotransmitter release, a GRAB norepinephrine sensor (ssAAV-9/2-hSyn1-GRAB_NE1h-WPRE-hGHp(A), titer 7.2×10E12 vg/ml) was used^29^. Mice (12-13 weeks) underwent stereotaxic surgery (Kopf Instruments, USA) for injections of AAVs carrying genetically encoded calcium indicators or fluorescent sensors. Anesthesia was achieved with the intraperitoneal injection of ketamine/xylazine (100/20 mg/kg) diluted in 0.9% saline. Isoflurane at 1-3% was given through a face mask to maintain anesthesia. Once the skull was exposed and bregma and lambda were aligned, a hand-drill was used to insert a nano-syringe (World Precision Instruments, USA) to reach coordinates for vector delivery. AAVs were injected in the perifornical area of hypothalamus (PeF: AP −1.4; ML 0.6; DV −5.1), the lateral hypothalamus (LH: AP −1.4; ML 1; DV −5.1), the locus coeruleus (LC: AP −5.4; ML −0.85; DV −3.7), the paraventricular thalamus (PVT: AP −0.3; ML 0; DV −3.7), and the cingulate cortex (CING: AP +1.4; ML −0.3; DV −1.7). Each viral vector was injected at 300 nl with a speed rate of 50 nl/min. Injected mice were group-housed for three weeks to allow the viruses to achieve proper diffusion and expression.

### Implantation of optic fibers and EEG/EMG electrodes

Three weeks after injections, mice underwent a second stereotaxic surgery to implant optic fibers and EEG along with EMG electrodes. Optic fibers (200 μm, 0.37 NA, glass and ferrules from Thorlabs) were inserted slowly above the same coordinates as the injected sites and secured to the skull with dental cement. Two gold plated screws were placed on the right hemisphere at coordinates (AP −2; ML +2.5) for EEG and (AP −6; ML +1.8) for the ground electrode. Two plastic-covered gold wires were inserted into the neck muscles and served as EMG electrodes. All fibers, electrodes and wires were fixed using the dental resin (Relyx 3M). The EEG/ground screws and EMG wires were soldered to a digital interface board (Pinnacle Technology INC, USA), and the whole implant was covered by dental cement (3M). Mice were allowed to recover from surgery for one week before the beginning of recordings.

### *In vivo* multisite fiber photometry

Animals were habituated to the recording cable and optical fibers in their open-top home cages for 3 days and kept tethered for the duration of the experiments. All recordings were performed between 16-18 weeks of age. To monitor *in vivo* changes in neuronal population spiking activity of specific cell types (calcium signals) or neurotransmitters (GRAB sensor signals) in multiple nuclei, a multi-channel fiber photometry system with two excitation wavelengths of 465 nm and 405 nm was used (bundle-imaging fiber photometry system, Doric Lenses). Interleaved excitation lights were sent into low-autofluorescence optic fiber patch cords (200 µm core, 0.37 NA, Doric Lenses), which were connected to the implanted fibers contained in a 1.25 mm diameter ferrule via a bronze sleeve. The GCaMP/GRAB emission fluorescence signals were collected through the same patch cords with the acquisition rate of 20 Hz using the Doric Neuroscience Studio software (Doric Lenses). The power of 465 nm and 405 nm lights at the tip of the optic fibers was set to 30-40 µW. The EEG/EMG signals were collected at the frequency of 2 kHz using the Pinnacle sleep recording systems and Sirenia software (Pinnacle Technology, USA). Recorded photometry traces were synchronized with EEG/EMG signals using the generated TTL pulses by the Doric Neuroscience Studio, which were recorded by the Sirenia software through a BNC cable.

### Immunofluorescence and histological confirmation

Mice were deeply anesthetized with sodium pentobarbital (150 mg/kg) and transcardially perfused with PBS, followed by 4% paraformaldehyde. Brains were removed, fixed in 4% paraformaldehyde and stored in PBS/sodium azide (0.02%) at 4 °C. Brains were later cut in coronal sections of 60 μm with a vibratome (VT1200S, Leica, Germany). Sections containing areas of viral injection were selected for immunofluorescence staining and sequentially incubated with the primary and secondary antibodies. The primary antibodies were mouse anti-orexin-A (1:200; sc-80263, Santa Cruz Biotechnology), rabbit anti-Cre (1:200; 69050, EMD Millipore), and rabbit anti-MCH (1:200; H-070-47, Phoenix Pharmaceuticals, Inc.). Images were acquired using the LSM 900 confocal microscope (Carl Zeiss, Germany) and analyzed using the Zen Blue software (Carl Zeiss, Germany). Microscopy investigation was also performed to locate the sites of optical fiber implantation. Data were excluded if the locations were not confirmed.

### Scoring and quantification of vigilance states

Scoring of vigilance states was performed manually using visual inspection of EEG/EMG recordings in 1-sec epochs, to allow precise scoring of microarousals and transition times, as described previously^30^. A custom toolbox written in MATLAB was used to visualize time and frequency characteristics of EEG/EMG traces. Wake was defined as periods of either theta band EEG activity accompanied by EMG bursts of movement-related activity, or periods that mice were immobile including feeding and grooming behaviors. NREMS was scored as periods with a relatively high amplitude low frequency delta band EEG activity accompanied by reduced muscle tone relative to wakefulness and associated with behavioral quiescence. REMS episodes were scored as sustained periods of theta band EEG activity and behavioral quiescence associated with muscle atonia, except for brief phasic muscle twitches. Transition to wake was defined as the first epoch with a rapid increase in muscle tone concurrent with low-amplitude fast-frequency EEG activity. Wake to NREMS transition was defined as the first epoch containing high-amplitude delta band activity appearing after EMG silencing. REMS onset was defined as an epoch with the absence of EMG tone concomitant with recursive synchronized theta rhythm. Number, duration, and state fragmentation of wakefulness, NREMS, and REMS were quantified using MATLAB scripts.

### Analysis of fiber photometry recordings

The output signals of the fiber photometry system were converted to *ΔF/F* = (*F* -*Fmean*) / *Fmean*, after amplification, digitization, and low-pass filtering. Dynamics of calcium signals and neurotransmitter release during vigilance state transitions were calculated by averaging the ΔF/F signal across wake-NREMS, NREMS-wake, NREMS-REMS, and REMS-wake transitions. Transition heatmaps were obtained by aligning the normalized *ΔF/F* signals (z-scored) at the transition points. To detect prominent activity in calcium signals, the *ΔF/F* signal was lowpass filtered using finite impulse response filters (0.2 Hz, 100-th order), and peaks were detected automatically using the MATLAB “findpeaks” function. For proper calculation of the density and interval of LC^NA^ reactivations during NREMS, only epochs with at least 3 reactivations were considered. Correlation between cell-type specific neuronal dynamics and sleep oscillations were measured using concurrent analysis of fiber photometry recordings and time-frequency analysis of EEG signals across NREMS and REMS and during transitions to these states.

### Spectral and time-frequency analysis

Raw electrophysiological recordings were down-sampled to 1000 Hz after applying a low pass filter (Chebyshev Type I, order 8, low pass edge frequency of 400 Hz, passband ripple of 0.05 dB). State-specific power spectral densities (PSDs) of EEG signals were calculated using the Welch’s method (MATLAB “pwelch” function), with 4-s windows having 50% overlap and 0.5-Hz frequency resolution. To correct for differences in absolute EEG power between animals, PSDs of each vigilance state was normalized to the weighted total PSD power across 1-45 Hz band of all vigilance states for each animal. The weighted total PSD power for each animal was calculated so that each state contributes equally to the total EEG power as described previously^31^. Time-frequency heatmaps were obtained using the multitaper spectral analysis.

### Detection of sleep spindles

NREMS spindles were detected automatically using an optimized wavelet-based method as described previously^30^. Briefly, the power of EEG signals within 9-16 Hz was estimated using the complex B-spline wavelet function, and smoothed using a 200 ms Hanning window, and then a threshold equal to 3 standard deviations above the mean was applied to detect the potential spindle events. Events shorter than 400 ms or longer than 2 s were discarded. Using band pass-filtered EEG signal in the spindle range (9–16 Hz), we automatically counted the number of cycles of each detected event and excluded those with <5 cycles or more than 30 cycles. To discard artefacts, events with a power in the spindle band lower than 6–8.5 Hz or 16.5– 20 Hz power bands were discarded.

### Phase-amplitude cross-frequency coupling

Correlation between phase of low frequency oscillations and fast frequency bands, i.e. phase-amplitude cross-frequency coupling, were assessed using the Modulation Index^32,33^. EEG/LFP signals were bandpass-filtered into theta (5-12 Hz), gamma (54-90 Hz), and fast-gamma (110-160 Hz) using finite impulse response filters with an order equal to three cycles of the low-cutoff frequency in both forward and reverse directions to eliminate phase distortion. The instantaneous theta phase and the gamma envelope were estimated using the Hilbert transform. Theta phase was discretized into 18 equal bins (*N*=18, each 20°) and the average value of gamma envelope within each bin was calculated. The resulting phase-amplitude histogram (*P*) was compared with a uniform distribution (*U*) using the Kullback-Leibler distance, 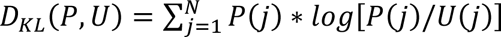, and normalized by *log(N)* to obtain the modulation index, *MI = D_KL_ / log(N)*. To explore other possible cross-frequency coupling patterns between different pairs of low and high-frequency bands, the comodulogram analysis was used^34^. Different frequency bands for phase (1-20 Hz, 1 Hz increments, 2 Hz bandwidth) and amplitude (20-200 Hz, 10 Hz increments, 20 Hz bandwidth) were used and MI values were calculated for all these pairs to obtain the comodulogram graph.

### Detection of phasic REMS events

Phasic REMS events, which are transient increase in theta power/frequency during REMS, were detected as described previously^35^. Briefly, EEG signal was bandpass-filtered between 4 and 12 Hz using finite impulse response filters with an order equal to three cycles of the low cutoff frequency, and the individual theta peaks were detected from the filtered signal. The interpeak interval time-series were smoothed using an 11-sample moving average window, and the smoothed interpeak intervals shorter than the 10th percentile were selected as candidate phasic REMS events. The candidate events with the following criteria were considered as phasic REMS: (1) minimum event duration of 900 ms; (2) minimum smoothed interpeak interval shorter than 5th percentile; (3) mean amplitude of theta peaks larger than mean amplitude of theta peaks across all REMS.

### Statistics

Animals of all experiments were randomly distributed for recordings, and individuals involved in vigilance state scoring were blinded to animals’ genotypes. The GraphPad Prism software was used to perform statistical analysis and compare different conditions via one-way or two-way ANOVA testes followed by multiple comparisons test, or two-sided *t*-tests for parametric data. All results are presented as mean ± SEM.

## Code availability

Custom-written MATLAB scripts for data analysis used in this study are available from the corresponding author upon request.

## Data availability

The datasets acquired for this study are available from the corresponding author upon request.

## Acknowledgements.

We thank Anne-Catherine Thomas for her help in animal genotyping. This work was supported by the Swiss National Science Foundation (grants 190605 to M.B., 201235 to M.T., and 31003A_182613 to A.V.) and the Novartis Foundation for Medical-Biological Research (grant 21B109 to M.B.).

## Author Contribution

G.B., G.C. and P.B. performed the experiments; M.B. and G.B. analyzed the data. M.B., G.B., M.T. and A.V. wrote the manuscript; All authors discussed the results and commented on the manuscript; M.B. and M.T. conceived and supervised the study.

## Supplementary figures

**Supp. Fig. 1:**
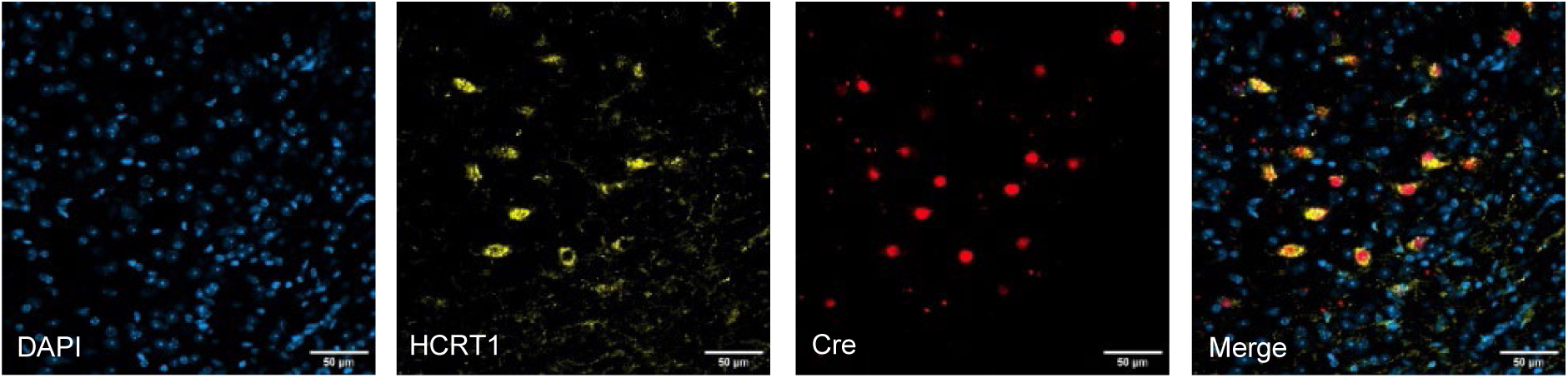
Validation of the *Hcrt-IRES-Cre* mouse line. Representative confocal acquisitions of a hypothalamic coronal section showing DAPI-, HCRT1-(Orexin-A), and Cre-stained neurons. Acquisitions merged show colocalization of HCRT1 and Cre. Scale bar: 50 μm.

**Supp. Fig. 2:**
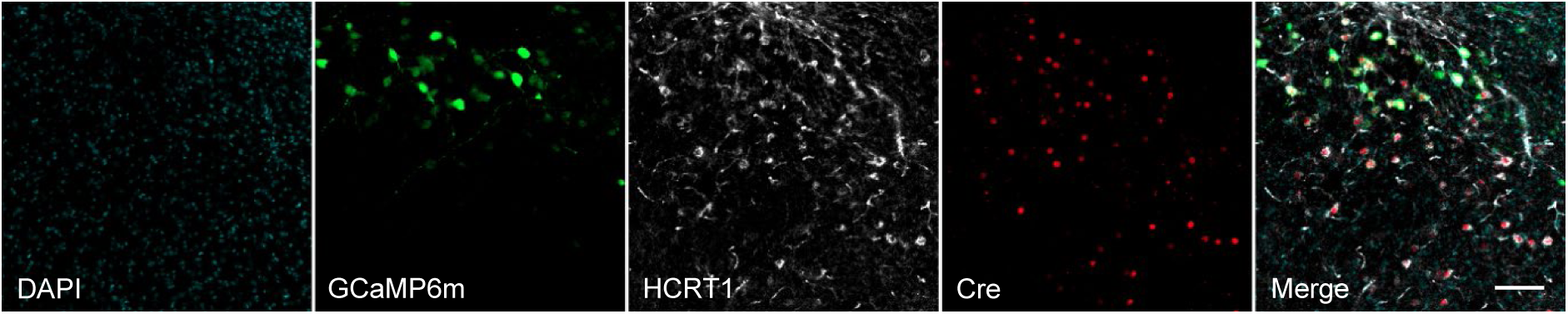
Validation of GCaMP6m expression in hypocretin neurons. Representative confocal acquisitions of a hypothalamic coronal section showing DAPI-, GCaMP6m-, HCRT1-(Orexin-A), and Cre-stained neurons. Acquisitions merged show colocalization of GCaMP6m-positive neurons and HCRT1/Cre neurons. Scale bar: 100 μm.

**Supp. Fig. 3:**
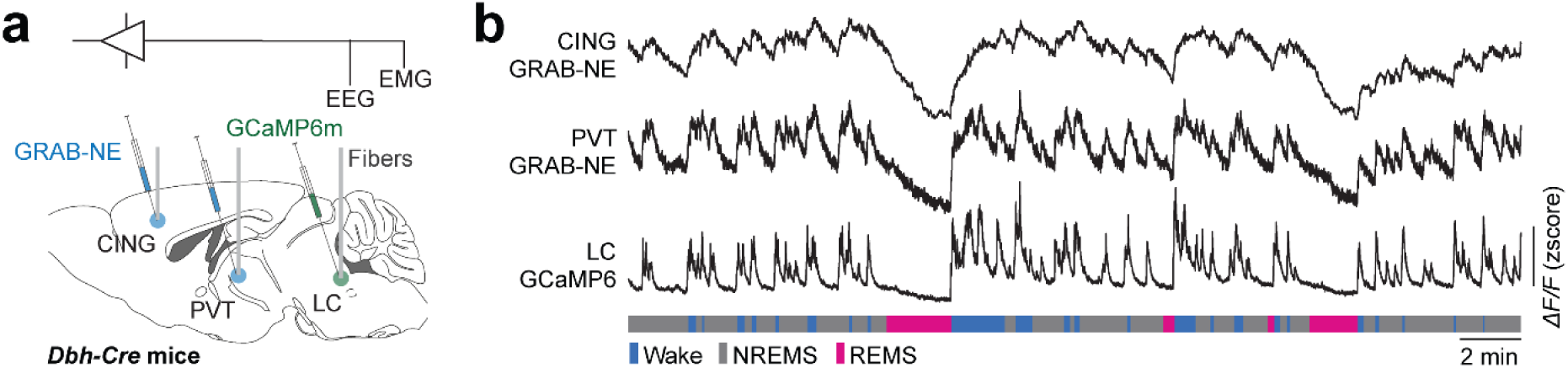
Locus coeruleus noradrenergic activity correlates with the release of norepinephrine in the midline thalamus and prefrontal cortex. **a)** Schematic of multisite fiber photometry of the locus coeruleus (LC), paraventricular thalamus (PVT), and cingulate cortex (CING). **b)** Multisite fiber photometry of LC noradrenergic activity and norepinephrine release in the midline thalamic and cortical regions in *Dbh-Cre* mice showed high synchronization between these signals across the vigilance states and their transitions. From bottom to top, hypnogram of vigilance states, *ΔF/F* of GCaMP6m in the LC, GRAB_NE_ signal in the PVT, and GRAB_NE_ signal in the CING.

**Supp. Fig. 4:**
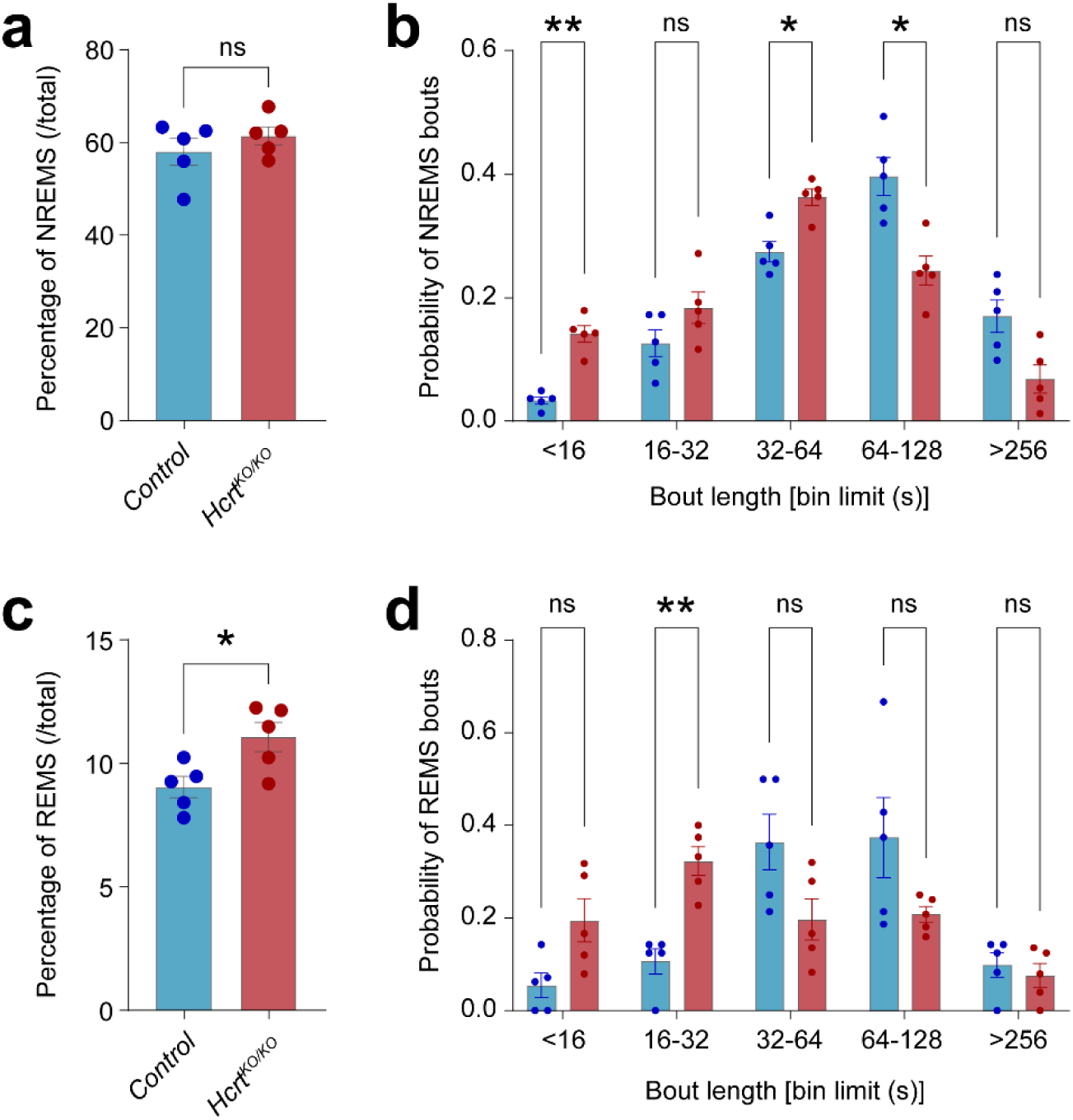
Quantification of sleep bouts between *Hcrt^KO/KO^* and control mice. **a)** Percentage of NREMS during the whole recording session (*p* = 0.3606, unpaired *t*-test). **b)** Distribution of NREMS bout durations indicates significantly shorter bouts in *Hcrt^KO/KO^* mice compared to controls (two-way ANOVA, interaction: *p* < 0.0001, followed by Sidak test). **c)** Percentage of REMS during the whole recording session (*p* = 0.0245, unpaired *t*-test). **d)** Distribution of REMS bout durations indicates significantly shorter bouts in *Hcrt^KO/KO^* mice compared to controls (two-way ANOVA, interaction: *p* = 0.0007, followed by Sidak test). n = 5 mice per group.

**Supp. Fig. 5:**
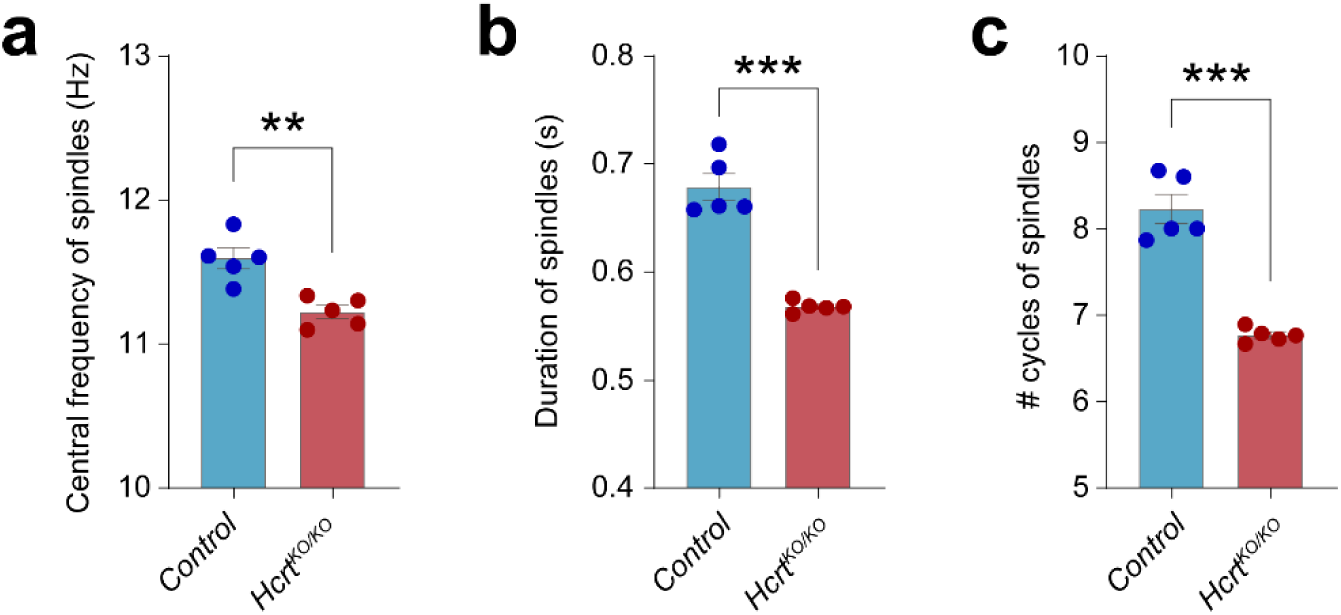
Characterization of NREMS spindles in *Hcrt^KO/KO^* and control mice. **a)** Central frequency of NREMS spindles is lower in *Hcrt^KO/KO^* mice compared to controls (*p* = 0.0024, unpaired *t*-test). **b)** Duration of NREMS spindles is shorter in *Hcrt^KO/KO^* mice compared to controls (*p* < 0.0001, unpaired *t*-test). **c)** Number of cycles per spindle is lower in *Hcrt^KO/KO^* mice compared to controls (*p* < 0.0001, unpaired *t*-test). n = 5 mice per group.

**Supp. Fig. 6:**
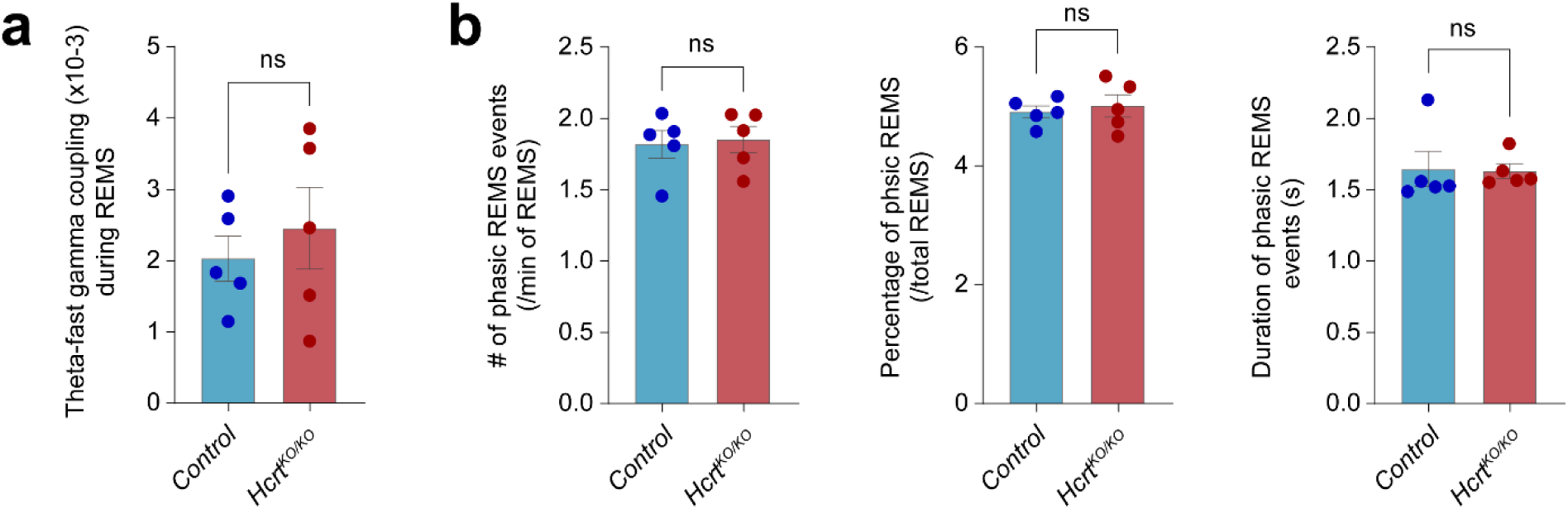
Characterization of theta-fast gamma coupling and phasic REMS events in *Hcrt^KO/KO^* and control mice. **a)** Theta-fast gamma coupling during REMS is not affected in the absence of hypocretin compared to controls (*p* = 0.5363, unpaired *t*-test). **b**) No significant differences were observed in the number (*p* = 0.8173, unpaired *t*-test), percentage (*p* = 0.6562, unpaired *t*-test) and duration (*p* = 0.9159, unpaired *t*-test) of phasic REMS events between *Hcrt^KO/KO^* and control mice. n = 5 mice per group.

## References

1. Scammell, T.E., Arrigoni, E. & Lipton, J.O. Neural Circuitry of Wakefulness and Sleep. Neuron 93, 747–765 (2017).

2. Lu, J., Sherman, D., Devor, M. & Saper, C.B. A putative flip-flop switch for control of REM sleep. Nature 441, 589–594 (2006).

3. Saper, C.B., Fuller, P.M., Pedersen, N.P., Lu, J. & Scammell, T.E. Sleep state switching. Neuron 68, 1023–1042 (2010).

4. Carter, M.E., et al. Mechanism for Hypocretin-mediated sleep-to-wake transitions. Proc Natl Acad Sci U S A 109, E2635–2644 (2012).

5. Mahler, S.V., Moorman, D.E., Smith, R.J., James, M.H. & Aston-Jones, G. Motivational activation: a unifying hypothesis of orexin/hypocretin function. Nat Neurosci 17, 1298–1303 (2014).

6. Sakurai, T. The role of orexin in motivated behaviours. Nat Rev Neurosci 15, 719–731 (2014).

7. de Lecea, L., et al. The hypocretins: hypothalamus-specific peptides with neuroexcitatory activity. Proc Natl Acad Sci U S A 95, 322–327 (1998).

8. Sakurai, T., et al. Orexins and orexin receptors: a family of hypothalamic neuropeptides and G protein-coupled receptors that regulate feeding behavior. Cell 92, 573–585 (1998).

9. Peyron, C., et al. Neurons containing hypocretin (orexin) project to multiple neuronal systems. The Journal of neuroscience : the official journal of the Society for Neuroscience 18, 9996–10015 (1998).

10. Nishino, S., Ripley, B., Overeem, S., Lammers, G.J. & Mignot, E. Hypocretin (orexin) deficiency in human narcolepsy. Lancet 355, 39–40 (2000).

11. Lin, L., et al. The sleep disorder canine narcolepsy is caused by a mutation in the hypocretin (orexin) receptor 2 gene. Cell 98, 365–376 (1999).

12. Chemelli, R.M., et al. Narcolepsy in orexin knockout mice: molecular genetics of sleep regulation. Cell 98, 437–451 (1999).

13. Osorio-Forero, A., et al. Noradrenergic circuit control of non-REM sleep substates. Curr Biol 31, 5009–5023 e5007 (2021).

14. Kjaerby, C., et al. Memory-enhancing properties of sleep depend on the oscillatory amplitude of norepinephrine. Nat Neurosci 25, 1059–1070 (2022).

15. Adamantidis, A.R., Zhang, F., Aravanis, A.M., Deisseroth, K. & de Lecea, L. Neural substrates of awakening probed with optogenetic control of hypocretin neurons. Nature 450, 420–424 (2007).

16. Carter, M.E., et al. Tuning arousal with optogenetic modulation of locus coeruleus neurons. Nat Neurosci 13, 1526–1533 (2010).

17. Saper, C.B., Scammell, T.E. & Lu, J. Hypothalamic regulation of sleep and circadian rhythms. Nature 437, 1257–1263 (2005).

18. Li, S.B., et al. Hyperexcitable arousal circuits drive sleep instability during aging. Science 375, eabh3021 (2022).

19. Seifinejad, A., Li, S., Possovre, M.L., Vassalli, A. & Tafti, M. Hypocretinergic interactions with the serotonergic system regulate REM sleep and cataplexy. Nat Commun 11, 6034 (2020).

20. Vassalli, A., et al. Electroencephalogram paroxysmal theta characterizes cataplexy in mice and children. Brain 136, 1592–1608 (2013).

21. Grujic, N., Tesmer, A., Bracey, E., Peleg-Raibstein, D. & Burdakov, D. Control and coding of pupil size by hypothalamic orexin neurons. Nat Neurosci 26, 1160–1164 (2023).

22. Ito, H., et al. Deficiency of orexin signaling during sleep is involved in abnormal REM sleep architecture in narcolepsy. Proc Natl Acad Sci U S A 120, e2301951120 (2023).

23. Lee, M.G., Hassani, O.K. & Jones, B.E. Discharge of identified orexin/hypocretin neurons across the sleep-waking cycle. The Journal of neuroscience : the official journal of the Society for Neuroscience 25, 6716–6720 (2005).

24. Mileykovskiy, B.Y., Kiyashchenko, L.I. & Siegel, J.M. Behavioral correlates of activity in identified hypocretin/orexin neurons. Neuron 46, 787–798 (2005).

25. Harris, G.C. & Aston-Jones, G. Arousal and reward: a dichotomy in orexin function. Trends Neurosci 29, 571–577 (2006).

26. Giardino, W.J., et al. Parallel circuits from the bed nuclei of stria terminalis to the lateral hypothalamus drive opposing emotional states. Nat Neurosci 21, 1084–1095 (2018).

27. Stanke, M., et al. Target-dependent specification of the neurotransmitter phenotype: cholinergic differentiation of sympathetic neurons is mediated in vivo by gp 130 signaling. Development 133, 141–150 (2006).

28. Parlato, R., Otto, C., Begus, Y., Stotz, S. & Schutz, G. Specific ablation of the transcription factor CREB in sympathetic neurons surprisingly protects against developmentally regulated apoptosis. Development 134, 1663–1670 (2007).

29. Feng, J., et al. A Genetically Encoded Fluorescent Sensor for Rapid and Specific In Vivo Detection of Norepinephrine. Neuron 102, 745–761 e748 (2019).

30. Bandarabadi, M., et al. A role for spindles in the onset of rapid eye movement sleep. Nat Commun 11, 5247 (2020).

31. Franken, P., Malafosse, A. & Tafti, M. Genetic variation in EEG activity during sleep in inbred mice. The American journal of physiology 275, R1127–1137 (1998).

32. Tort, A.B., Komorowski, R., Eichenbaum, H. & Kopell, N. Measuring phase-amplitude coupling between neuronal oscillations of different frequencies. Journal of neurophysiology 104, 1195–1210 (2010).

33. Bandarabadi, M., et al. Dynamic modulation of theta-gamma coupling during rapid eye movement sleep. Sleep 42, 1–11 (2019).

34. Tort, A.B., et al. Dynamic cross-frequency couplings of local field potential oscillations in rat striatum and hippocampus during performance of a T-maze task. Proc Natl Acad Sci U S A 105, 20517–20522 (2008).

35. Mizuseki, K., Diba, K., Pastalkova, E. & Buzsaki, G. Hippocampal CA1 pyramidal cells form functionally distinct sublayers. Nat Neurosci 14, 1174–1181 (2011).

